# Cell Programmed Nutrient Partitioning in the Tumor Microenvironment

**DOI:** 10.1101/2020.08.10.238428

**Authors:** BI Reinfeld, MZ Madden, MM Wolf, A Chytil, JE Bader, AR Patterson, AS Cohen, A Ali, BT Do, A Muir, CA Lewis, RA Hongo, KL Young, RE Brown, VM Todd, T Huffstater, A Abraham, RT O’Neil, MH Wilson, F Xin, MN Tantawy, WD Merryman, RW Johnson, CS Williams, EF Mason, FM Mason, KE Beckermann, MG Vander Heiden, HC Manning, JC Rathmell, WK Rathmell

**Author notes:** These authors contributed equally to this work (BI Reinfeld, MZ Madden, JC Rathmell, WK Rathmell).

## Abstract

The tumor microenvironment (TME) includes transformed cancer and infiltrating immune cells^1,2^. Cancer cells can consume large quantities of glucose through Warburg metabolism^3,4^ that can be visualized with positron emission tomography (PET). While infiltrating immune cells also rely on glucose, disruptions to metabolism can contribute to tumor immunological evasion^5–9^. How immune cell metabolism is programmed or restrained by competition with cancer cells for nutrients, remains uncertain. Here we used PET tracers to measure the accessibility of glucose and glutamine to cell subsets in the TME. Surprisingly, myeloid cells including macrophages were the greatest consumers of intra-tumoral glucose, followed by T cells and cancer cells. Cancer cells, in contrast, had the highest glutamine uptake. This distinct nutrient partitioning was programmed through selective mTORC1 signaling and glucose or glutamine-related gene expression. Inhibition of glutamine uptake enhanced glucose uptake across tumor resident cell types and shifted macrophage phenotype, demonstrating glucose is not limiting in the TME. Thus, cancer cells are not the only cells in tumors which exhibit high glucose uptake *in vivo* and instead preferentially utilize glutamine over other cell types. We observe that intrinsic cellular programs can play a major role in the use of some nutrients. Together, these data argue cell selective partitioning of glucose and glutamine can be exploited to develop therapies and imaging strategies to alter the metabolic programs of specific cell populations in the TME.

---

The founding observation in cancer metabolism was that sections of tumors produced significant lactate in the presence of oxygen^4^. These results gave rise to the theory that glycolysis in the presence of oxygen, also known as aerobic glycolysis or the Warburg effect, supports cancer cell growth and proliferation. Subsequently, the Warburg effect has been observed in other rapidly proliferating cells, including activated immune cells^5^, in which biosynthetic demands can be met by shunting glucose carbons into pathways to generate amino acids, nucleotides, and lipids^3^. Recent data from *in vivo* carbon labeling studies have confirmed that glucose supports anabolic metabolism in transformed cells^10^ and in T cells^11^. In addition, glutamine metabolism can support aerobic glycolysis as an anaplerotic or distinct fuel source yet can also restrain glucose-dependent inflammatory differentiation and function of both macrophages and T cells^12–15^. Glucose uptake in tumor cells is illustrated as a hallmark of malignancy by the prominent role of ^18^F-fluorodeoxyglucose positron emission tomography (FDG-PET) imaging in detecting cancers and monitoring therapeutic responses. Based on these metabolic programs, nutrient competition and depletion of glucose in the TME by cancer cells has been proposed as a metabolic mechanism of immunosuppression^8,9^. Recent publications, however, have measured high micromolar to millimolar glucose concentration in the TME^16–18^. Further, the metabolic phenotypes of T cells can persist even after removal from the TME^18^. The extent to which immune cell metabolism is programmed or dependent on nutrient competition between metabolically active cancer cells and immune cells remains uncertain. Here we analyzed the accessibility of glucose and glutamine to specific cell subsets in the TME and show that metabolites partition into distinct cellular compartments with programmed metabolisms that should be considered when developing anticancer therapies and PET imaging probes for cancer diagnosis and monitoring.

## Heterogeneous glucose uptake in the TME

Based on observations that both cancer and inflammatory conditions increase glucose uptake, we hypothesized that immune cells could contribute significantly to glucose consumption in the TME. To test this, we developed a method to measure *in vivo* nutrient availability and uptake across cancer and immune cell populations isolated from the heterogenous murine tumors. Steady state nutrient abundance was first measured in the TME in tissue interstitial fluid (IF) from freshly resected human clear cell renal cell carcinoma (ccRCC) specimens using mass spectrometry^16^ (**Fig 1a**). Glucose, glutamine, and lactate were all detectable in the TME; their concentrations were similar to, if not higher than, matched normal kidney tissue; and glucose levels were in a range similar to that found in plasma^19^. These data suggest that concentrations of these nutrients may not necessarily be limiting for different tumor cell populations. To quantify how much glucose is accessible by distinct cell populations in the TME, subcutaneous tumors were grown to a point clearly visualized by FDG-PET imaging (**Fig 1b**), and per cell *in vivo* FDG uptake was measured in magnetic bead-fractionated tumor cell subsets (**Fig 1c**). Using CD45 positive selection microbeads, we fractionated total tumor cells into highly enriched CD45^-^ cancer and stromal cell and CD45^+^ immune cell populations (**Fig 1d, Extended Data Fig 1a-c**). Unfractionated tumor cells demonstrated higher FDG avidity than control tissue splenocytes consistent with canonical Warburg metabolism in tumors compared to normal tissue. Strikingly, tumor CD45^+^ immune cells had significantly higher per cell FDG uptake than CD45^-^ cancer cells (**Fig 1e**). Tumor-infiltrating immune cells also had higher FDG avidity in CT26 (**Fig 1f)** and Renca (**Fig 1g**) subcutaneous tumors. Renca tumors grown orthotopically in the kidney demonstrated higher per cell FDG avidity in immune cells, suggesting tissue site independence (**Fig 1h**). Finally, we measured FDG uptake in the TME of two spontaneous carcinogenesis models. Infiltrating immune cells had higher FDG uptake compared to EPCAM^+^ cancer cells in both PyMT genetically engineered mouse model (GEMM) breast cancer tumors and azoxymethane/dextran sodium sulfate-induced (AOM/DSS) inflammatory colon cancer tumors. (**Fig 1i, Extended Data Fig 1d-f**). Together, these results show that glucose is available in the TME and preferentially partitions into infiltrating immune cells more so than cancer cells across multiple models.

**Fig. 1.**
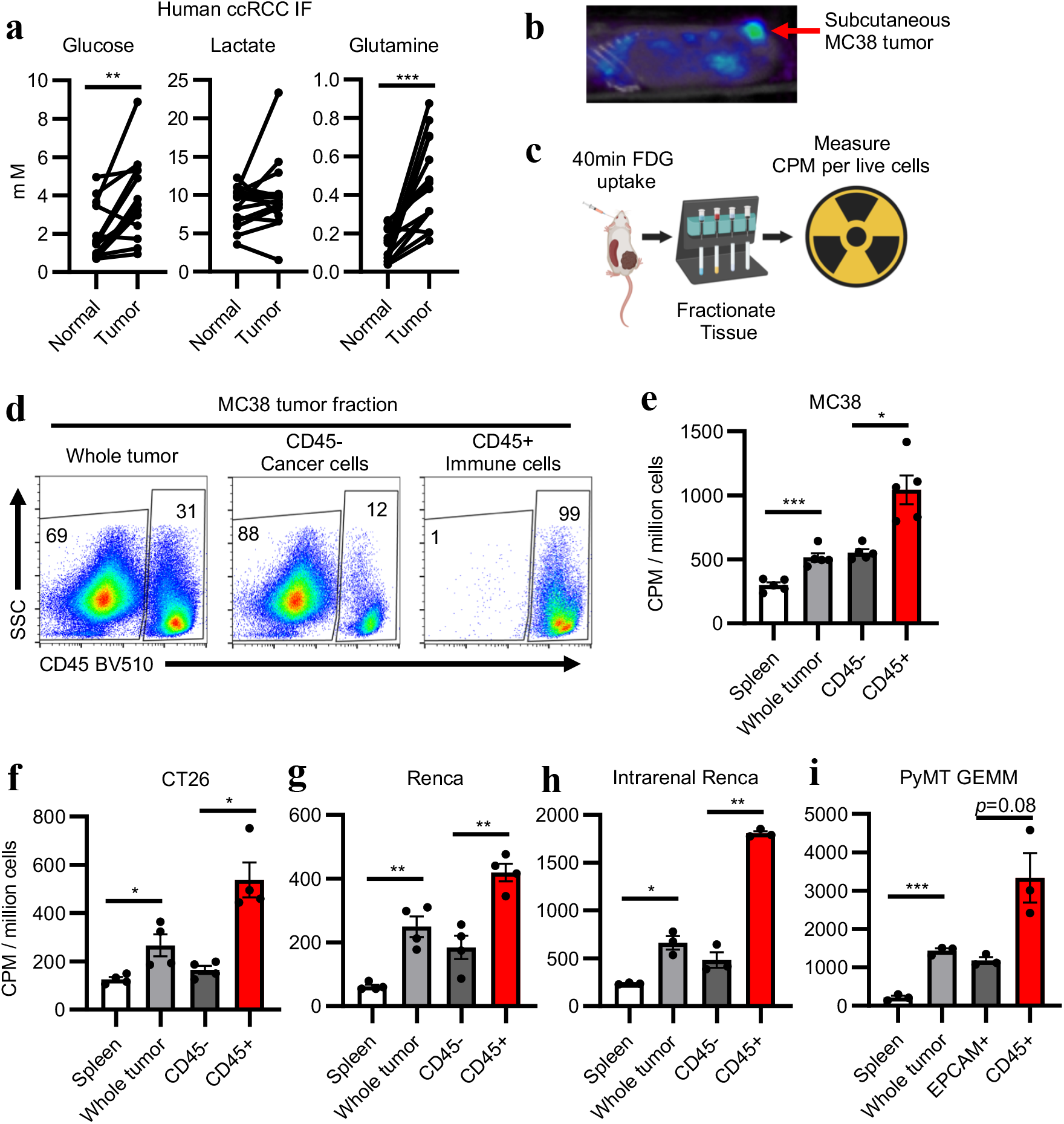
Glucose is present in the TME and is preferentially consumed by immune cells. **a**, Quantification of IF metabolites from patient ccRCC tumor and matched adjacent normal kidney tissue (n=14 patients). **b**, FDG PET-CT image of subcutaneous MC38 tumor. **c**, Experimental schema for isolation and characterization of FDG-avid tumor cell populations. **d**, Representative flow cytometry analysis of MC38 whole tumor, CD45+ immune cell, and CD45-cancer cell fractions gated on live cells. **e-i**, Per cell FDG avidity in designated tumor cell fractions in subcutaneous MC38 (n=5 mice) (**e**), CT26 (n=4 mice) (**f**), and Renca (n=4 mice) (**g**) tumors; intrarenal Renca tumors (n=3 mice) (**h**); and PyMT GEMM tumors (n=3 mice) (**i**). Each data point represents a biological replicate and error bars are SEM. e-i are data from representative studies performed independently at least twice. P values were calculated using paired 2-tailed t-test for (a) and Welch’s 2-tailed t-test for (e-i). \**p*<0.05, ** *p*<0.01, *** *p*<0.001. ccRCC: clear cell renal cell carcinoma; CPM: counts per minute; FDG PET-CT: 18-fluorodeoxyglucose positron emission tomography computed tomography; GEMM: genetically engineered mouse model; IF: interstitial fluid; PyMT: Polyoma virus middle T antigen; TME: tumor microenvironment.

We validated that this approach measures *in vivo* per cell glucose uptake using multiple approaches. The FDG uptake phenotypes were independent of sample viability and cell yield across biological replicates and across tumor models (**Extended Data Fig 1b-c**). The dynamic range of ^18^F-FDG uptake was found to have a multiple-log scale of linearity (**Extended Data Fig 2a**). Immune cells obtained from MC38 tumors demonstrated low intravenous labeling of CD45, supporting that fractionated tumor immune cells were tumor-resident (**Extended Data Fig 2b**). To confirm that measured FDG uptake occurred *in vivo* and was not an artifact of tumor processing, we incubated MC38 FDG-naïve tumor cell suspensions with FDG-treated tumor digest supernatant and found that *ex vivo* FDG uptake accounted for only a small fraction of the final FDG signal (**Extended Data Fig 2c**). Finally, to test if CD45 positive selection artificially increased *ex vivo* FDG uptake by immune cells, we generated Thy1.1^+^ MC38 cells and isolated cancer cells using Thy1.1 positive selection microbeads. Consistent with data generated with CD45 microbeads, Thy1.1^-^ immune cells demonstrated higher FDG avidity than Thy1.1^+^ cancer cells, indicating that magnetic purification via CD45 microbeads does not artifactually increase FDG uptake by immune cells (**Extended Data Fig 2d-e**).

## TME myeloid cells uptake more glucose than other cell types evaluated

We next sought to identify which immune cells in the TME have the greatest access and ability to uptake glucose. Tumor infiltrating immune cells are comprised of multiple hematopoietic lineages^1,2^. CD3^+^ T cells are a primary focus of modern cancer immunotherapy, and T cell activation and infiltration are associated with elevated glycolysis^7,20–22^. CD11b^+^ myeloid cells can play anti- and pro-tumor roles in the TME and are also being actively investigated as a target for immunotherapy^23^. Flow cytometry characterization of immune infiltrates in various TMEs demonstrated extensive heterogeneity in immune cell composition (**Extended Data Fig 3a-f**). Given the rich T cell and myeloid infiltrate in MC38 tumors (**Fig 2a**) and high cell viability and yield, we first sought to compare the FDG uptake between infiltrating tumor T cells, myeloid cells, and cancer cells in MC38. T cells were isolated using CD4/8 positive selection beads. Tumor-associated T cells had greater *in vivo* FDG avidity than resting splenic T cells and had similar ability to uptake glucose as cancer cells (**Fig 2b-c, Extended Data Fig 4a**), suggesting that T cells are not deprived or out-competed for glucose in the TME. T cell glucose uptake was significantly lower, however, than that of the remaining CD45^+^ non-T cells.

**Fig. 2.**
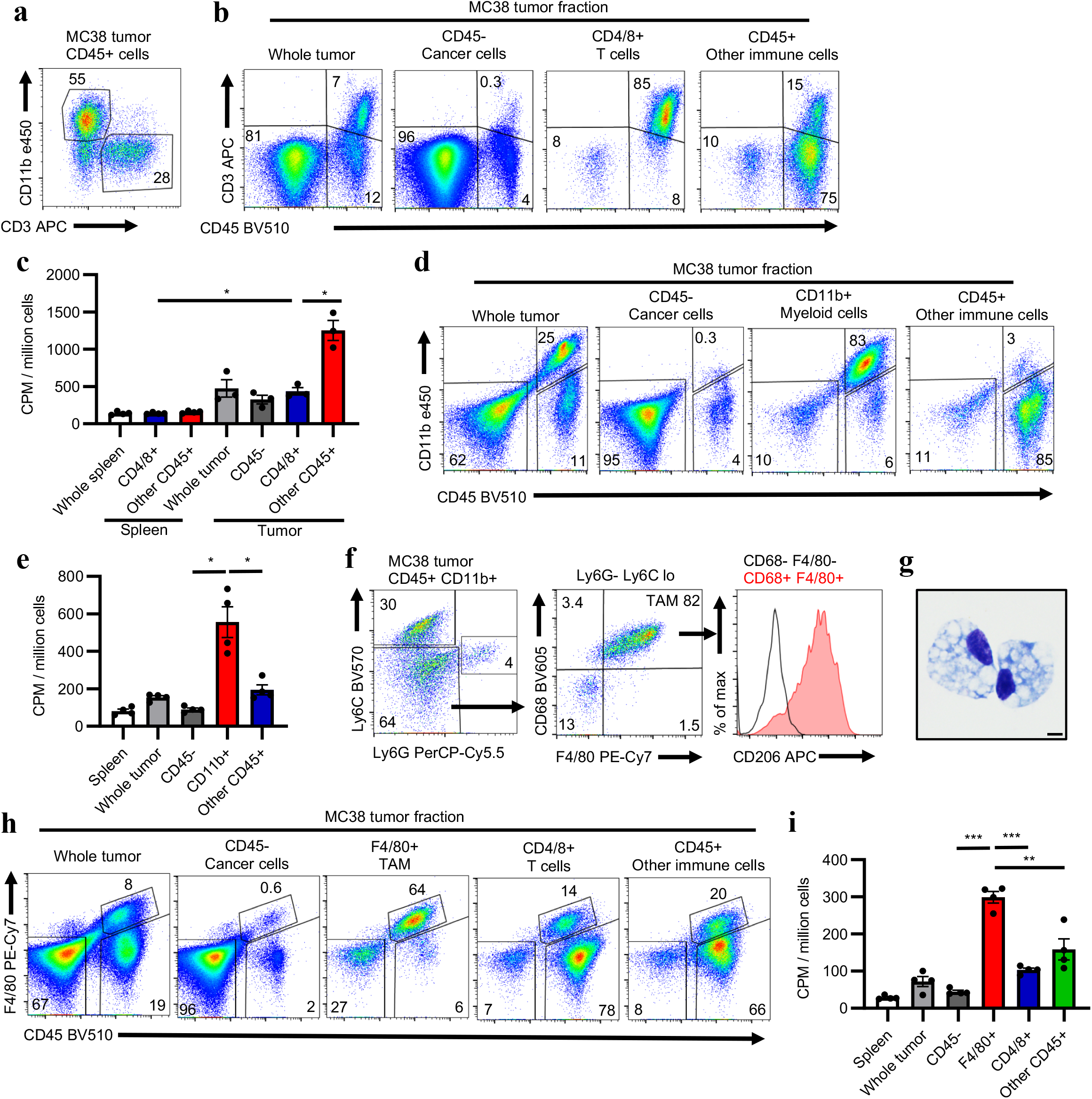
Myeloid cells uptake the most glucose in the TME. **a**, Representative flow cytometry plot from MC38 tumor gated on CD45+. **b**, Representative flow cytometry plots gated on live cells of indicated MC38 tumor cell fractions with T cell isolation using anti-CD4/8 microbeads. **c**, Per cell FDG avidity in designated spleen and MC38 tumor cell fractions using anti-CD4/8 microbeads (n=3 mice). **d**, Representative flow cytometry plots gated on live cells of indicated MC38 tumor cell fractions with myeloid cell isolation using anti-CD11b microbeads. **e**, Per cell FDG avidity in designated MC38 tumor cell fractions using anti-CD11b microbeads (n=4 mice). **f**, Flow cytometry characterization of TAM. **g**, Representative micrograph of anti-F4/80 microbead-isolated TAM, scale bar = 5μm. **h**, Representative flow cytometry plots gated on live cells of indicated MC38 tumor cell fractions with TAM isolation using anti-F4/80 microbeads. **i**, Per cell FDG avidity in designated MC38 tumor cell fractions using anti-F4/80 microbeads (n=4 mice). Each data point represents a biological replicate and error bars are SEM. Representative studies were performed independently at least twice. P values were calculated using Welch’s 2tailed t-test. * *p*<0.05, ** *p*<0.01, *** *p*<0.001. CPM: counts per minute; FDG: 18-fluorodeoxyglucose; TAM: tumor-associated macrophage; TME: tumor microenvironment.

To characterize the non-T cell CD45^+^ cells we performed a reciprocal experiment isolating myeloid cells using CD11b positive selection beads (**Fig 2d-e, Extended Data Fig 4b**). CD11b^+^ myeloid cells did indeed have higher FDG uptake than cancer cells and other immune cells in MC38 tumors. CT26 tumors showed a similar phenotype (**Extended Data Fig 4c-f**). Flow cytometry analysis of CD45^+^ CD11b^+^ cells from MC38 tumors demonstrated a high proportion of Ly6G^-^, Ly6C^lo^, F4/80^hi^, CD68^+^, CD206^hi^ cells consistent with tumor associated macrophages (TAM) (**Fig 2f**). Isolated F4/80^hi^ cells had histiocytic morphology (**Fig 2g**), consistent with TAM classification and TAM isolated using F4/80 positive selection microbeads demonstrated the highest FDG uptake of any MC38 tumor cell population (**Fig 2h-i, Extended Data Fig 4g**). These studies support a model where myeloid cells including TAM consume the most glucose in the TME. Consistent with our findings, myeloid infiltration is known to drive FDG avidity in nontumor bearing lymph nodes in human and mouse gynecological malignancies^24^. Our data extend these findings directly to the solid TME.

## TME myeloid cells are highly metabolically active

To further understand the metabolic phenotypes of distinct tumor cell populations and establish if these were cell-intrinsic and persistent or dependent on the TME, we conducted extracellular flux assays on microbead-fractionated MC38 tumor cells (**Fig 3a-d**). Isolated F4/80^+^ TAM maintained higher basal cellular extracellular acidification rate (ECAR) and basal and maximal mitochondrial oxygen consumption rate (OCR) than tumor infiltrating T cells and cancer cells, consistent with higher TAM glycolytic and oxidative metabolism. Next, we transcriptionally profiled MC38 tumor flow-sorted cell populations including CD45^-^ cancer cells, CD11b^+^ F4/80^hi^ TAM, CD11b^+^ F4/80^lo^ myeloid cells, CD3^+^ CD8a^+^ T cells, and CD3^+^ CD8a^-^ (CD4^+^) T cells (**Extended Data Fig 5a-b**). Unsupervised clustering of differentially expressed metabolism-related transcripts grouped samples by cell identity (**Fig 3e**). Glucose transporters demonstrated population-specific expression, with cancer cells and myeloid cells expressing high levels of *Slc2a1* (GLUT1) and T cells expressing high levels of *Slc2a3* (GLUT3) (**Fig 3f, Extended Data Figs 5c-d**). Hexokinase isoforms *Hk2* and *Hk3,* which catalyze glucose phosphorylation in cells as the rate-limiting initial phosphorylation of glucose in glycolysis, are specifically expressed in myeloid cells in contrast to broadly expressed *Hk1* (**Fig 3g**). Anabolic metabolism is supported by mTORC1 signaling. Flow cytometry analysis demonstrated high levels of phosphorylated ribosomal protein S6 downstream of mTORC1 in myeloid and cancer cells of MC38, CT26, and human ccRCC tumors (**Fig 3h-k, Extended Data Fig 5e**). In combination, these data show that myeloid cells including TAM are programmed to exhibit a metabolic phenotype of high glucose uptake and are a metabolically active cell population in the TME.

**Fig. 3.**
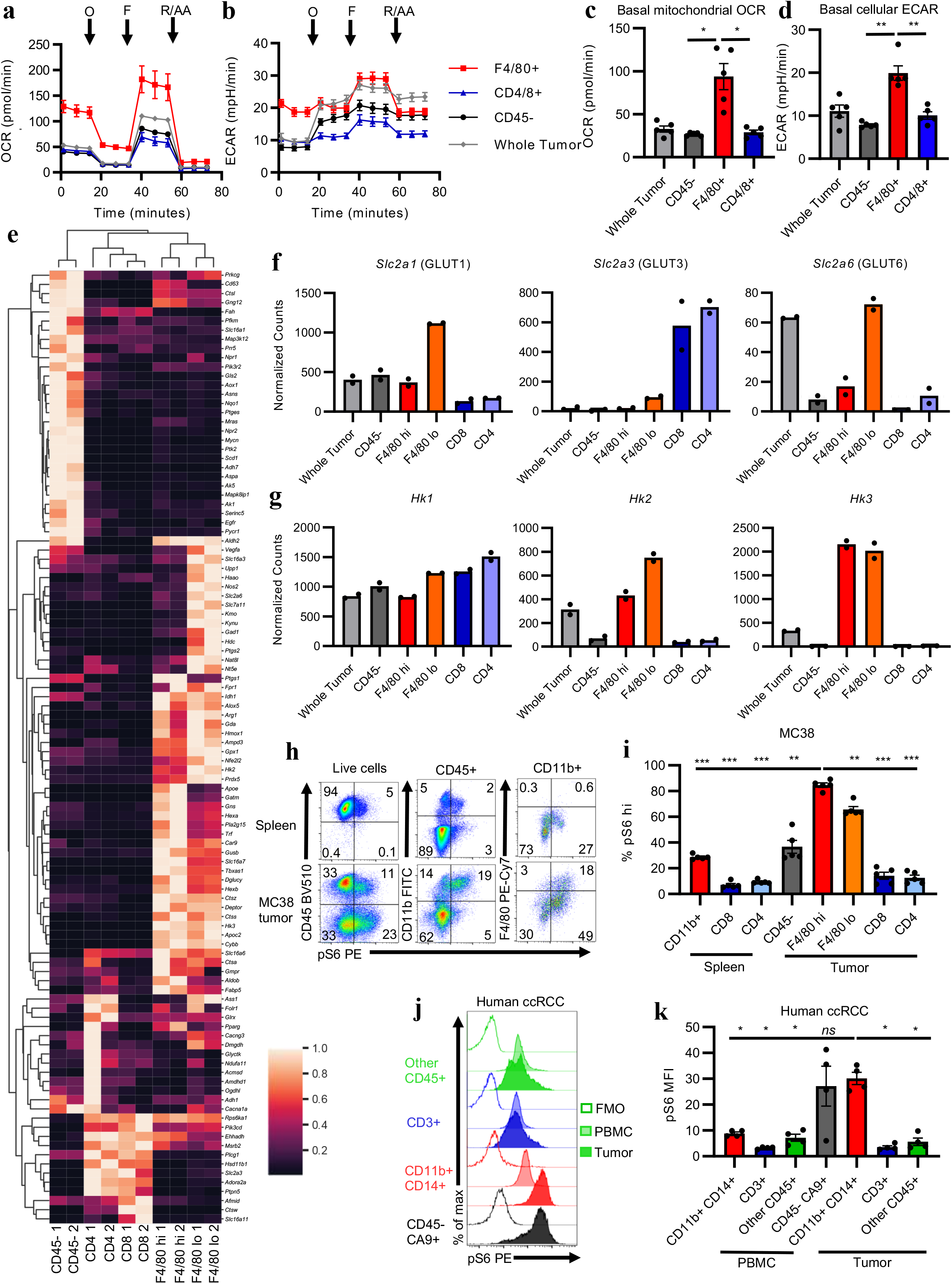
Characterization of tumor cell populations reveals regulators of glucose uptake. **a-b**, Representative OCR (**a**) and ECAR (**b**) tracings of MC38 tumor cell fractions with indicated injections of oligomycin (O), FCCP (F), and rotenone and antimycin A (R/AA). **c-d** Basal mitochondrial OCR (**c**) and cellular ECAR (**d**) of MC38 tumor fractions (n=5 mice). **e**, Unsupervised cluster analysis of differentially expressed metabolic mRNA transcripts of whole tumor, CD45-cancer cell, CD45+ CD11b+ F4/80hi TAM, CD45+ CD11b+ F4/80lo myeloid cell, CD45+ CD3+ CD8a+ T cell, and CD45+ CD3+ CD8a-(CD4+) T cell flow-sorted MC38 tumor populations (n=2 mice). **f-g**, Glucose transporter (**f**) and hexokinase (**g**) mRNA transcript levels of indicated MC38 tumor populations (n=2 mice). Dotted line approximates limit of detection. **hi**, Representative flow cytometry plots (**h**) and quantification (**i**) of pS6 levels in indicated MC38 tumor and spleen populations. **j-k**, Representative flow cytometry histograms (**j**) and quantification (**k**) of pS6 levels in cancer cells (CD45-CA9+), myeloid cells (CD45+ CD11b+ CD14+), T cells (CD45+ CD3+), and other immune cells (CD45+ CD3-CD14-) from patient ccRCC tumor and PBMC (n=4 patients). Each data point represents a biological replicate and error bars are SEM. **a-d** and **f-g** are representative of at least two independent experiments. P values were calculated using Welch’s 2-tailed t-test for (c-d) and Brown-Forsythe ANOVA test for (h, j). * *p*<0.05, ** *p*<0.01, *** *p*<0.001. ECAR: extracellular acidification rate; FCCP: Trifluoromethoxy carbonylcyanide phenylhydrazone; FMO: fluorescence minus one; MFI: median fluorescence intensity; OCR: oxygen consumption rate; PBMC: peripheral blood mononuclear cells; pS6: phosphorylated ribosomal protein S6 (Ser235/236).

## Glutamine partitions into cancer cells

In addition to glycolysis, some cells utilize glutamine to maintain mitochondrial metabolites for ATP and macromolecule synthesis^25,26^. Recent work has shown that inhibiting TME glutamine metabolism can impair cancer cell growth while increasing anti-tumor immunity^12,13,15^. In contrast to our finding that glucose uptake is highest in tumor-infiltrating immune cells, we hypothesized that glutamine uptake might more preferentially support cancer cell metabolism. Studies have uncovered that malignant cell MYCN and ATF4 drive glutamine utilization across many tumor types^27–30^. In MC38 tumors, *Mycn* and *Atf4* were more highly expressed in cancer cells than immune cells (**Fig 4a**). Glutamine metabolism enzymes *Aspa*^31,32^, *Asns*^33^, and *Gls2*^34^ were also specifically expressed in the MC38 cancer cells (**Extended Data Fig 6a**). To test if glutamine was specifically taken up by cancer cells, we measured ^18^F-glutamine uptake in MC38, CT26, and Renca tumors. In contrast to glucose, glutamine preferentially accumulated in cancer cells as compared with immune cells across all three models (**Fig 4b-c, Extended Data Fig 6b-e)**.

**Fig. 4.**
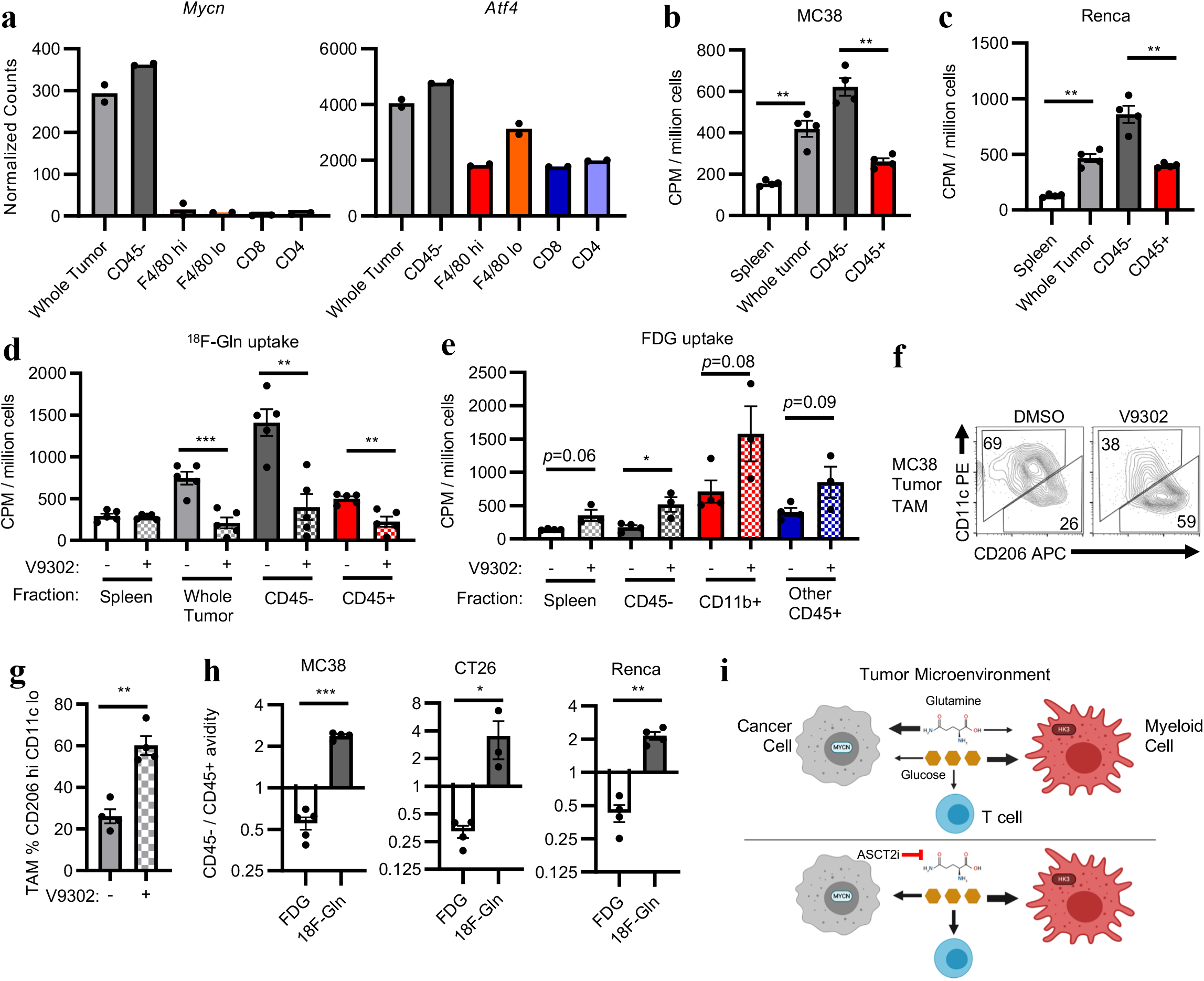
Glutamine partitions into cancer cells in the TME. **a**, Glutamine-related transcription factor mRNA transcript levels of indicated flow-sorted MC38 tumor cell populations (n=2 mice). Dotted line approximates limit of detection. **b-c**, Per cell 18F-Gln avidity in designated tumor cell fractions in subcutaneous MC38 (n=3-4 mice) (**b**) and Renca (n=4 mice) (**c**) tumors. **d,** Per cell 18F-Gln avidity in designated MC38 tumor cell fractions from mice treated with V9302 or DMSO (n=5 mice) **e**, Per cell FDG avidity in designated MC38 tumor cell fractions in subcutaneous MC38 tumors from mice treated with DMSO or V9302 (n=3-4 mice). **f**, Representative flow cytometry analysis gated on CD45+ CD11b+ Ly6G-Ly6Clo TAM from MC38 tumors in mice treated with DMSO or V9302 (n=4 mice). **g**, Quantification of macrophage populations from MC38 tumors in mice treated with DMSO or V9202 (n=4 mice). **h**, Ratio of CD45-divided by CD45+ per cell 18F-nutrient avidities in indicated tumors (n=3-5 mice). **i**, Model for nutrient partitioning in the TME. Each data point represents a biological replicate and error bars are SEM. **b-h** are representative of at least two independent experiments. P values were calculated using Welch’s 2-tailed t-test for a-c and e-h, one 1-tailed for d. * *p*<0.05, ** *p*<0.01, *** *p*<0.001. 18F-Gln: 18F-4-fluoroglutamine; CPM: counts per minute; TAM: tumor-associated macrophage; TME: tumor microenvironment; V9302: ASCT2 inhibitor.

Given our finding that glutamine uptake is highest in cancer cells, we tested a glutamine transport inhibitor, V9302^35,36^ to reprogram cell metabolism in the heterogenous TME. Glutamine transporters are expressed throughout all cell types in the TME (**Extended Data Fig 7a**). MC38 tumors from mice treated with V9302 trended toward a smaller size compared to vehicle-treated controls (**Extended Data Fig 7b**). Acute V9302 treatment decreased glutamine uptake by all cells in the TME (**Fig. 4d**). After 5 days of V9302 treatment, we observed increased glucose uptake in cancer cells, with a trend towards increased glucose uptake in splenocytes, CD11b^+^ myeloid and other CD45^+^ cells in the TME (**Fig 4e, Extended Data Fig 7c**), demonstrating that access to glutamine restrains glucose metabolism *in vivo* and that cells were able to access and increase glucose uptake beyond basal levels. Glutamine inhibition also increased tumor TAM abundance (**Extended Data Figure 7d**) and shifted macrophages to a more M2-like immunophenotype (**Fig 4f**) in line with the role of glutamine synthetase in these tumor suppressive cells^37^.

## Selective Nutrient Partitioning

Rather than cells depleting and consuming a common nutrient pool, specific cell types in the TME appeared to display programs for preferential nutrient utilization. Comparing the ratios of CD45-to CD45+ per-cell ^18^F-nutrient uptake, ^18^F-glutamine uptake was significantly higher than FDG uptake for CD45-cancer cells versus immune cells in MC38, CT26, and Renca tumors (**Fig 4g)**, suggests that ^18^F-glutamine is a more cancer-cell specific tracer than FDG in at least some tumors. Taken together, this work suggests that glutamine preferentially accumulates in cancer cells in the TME in contrast to glucose which is taken up by immune cells (**Fig. 4h**). These results also support our human tumor interstitial fluid from ccRCC findings that argue glucose is not limiting in the TME of many tumors^16–18^.

Our findings stand in contrast to the model that cancer cells are the only TME resident cells with high glucose uptake and thus contributing to immune cell dysfunction via nutrient competition. This work reveals that diverse cell populations can preferentially uptake different metabolites from a common pool in the TME in line with recent publications^38,39^. Previously, direct comparisons of *in vivo* glucose utilization between cancer cells and immune cells have not been reliably quantified ^9,40–42^. Our work demonstrates tumor myeloid cells consume significantly more glucose than tumor infiltrating T cells or cancer cells. This may have implications for a new generation of metabolism-targeting agents as well as myeloid targeting therapies, with the potential for therapeutics to either enhance or impair tumor-related inflammation. These data also support the notion that targeting glutamine metabolism presents a specific strategy to hamper cancer cell growth. Intriguingly, this approach also increases glucose consumption by all TME resident cell types and may alter antitumor immunity^12,13,15^. Given the current interest in glutamine-targeting therapeutics, our findings contribute a model of nutrient partitioning that supports their further development.

Previous studies have suggested competition for glucose in the TME between cancer cells and T cells contributes to immunosuppression^8,9,43^. While this may be the case in some cancers, our data show that for some tumors not only is glucose not limiting, but T cells and cancer cells have similar per cell glucose uptake and have the capacity to increase this level when glutamine uptake is restricted. Other groups have presented data that tumor hypoxia and oxidative metabolism may drive T cell dysfunction in the TME^44^. We find, however, that myeloid cells consume more glucose than other TME cells on a per cell basis. There is growing evidence that glycolytic cancer cell transcriptional programs are associated with immunosuppressive TMEs and directly recruit suppressive myeloid cells^45–47^ while TAM glycolysis can drive hypoxia via endothelial dysfunction^48^. Our work further supports a model where glycolytic tumors are immunoinhibitory not directly due to nutrient deficiencies but rather because of large scale microenvironmental changes, such as the presence of glycolytic myeloid cells and accumulation of waste products.

FDG imaging intensity is not relatable to *in vivo* ^13^C-glucose labeling^49^, and FDG PET in not suitable for imaging responses to immune checkpoint blockade using traditional assessments. Additionally, there are some tumor types like ccRCC that are not typically FDG-avid while harboring glycolysis-inducing mutations. In support of these clinical perplexities, our data question the notion that FDG tumor avidity is predominantly caused by cancer cell metabolism and growth. The FDG imaging features of a tumor may also be dependent on the type and activation state of a patient’s tumor immune infiltrate. Supporting this, recent work has shown increased FDG avidity in lung cancer is associated with more CD68^+^ TAM, which was interpreted as an increase in cancer cell glycolysis due to macrophage secretion of TNFα^42^. Additionally, qualitative assessments of FDG via tissue autoradiography noted that this tracer accumulated in the histiocytic periphery of mouse tumors.^50^ Our data support the model where TAM directly consume FDG and thus account for the measured tumor glucose uptake due to their own intrinsic glycolytic activity. This model also lends an explanation for regional variability in FDG avidity observed on PET imaging as well as the PET avid nature of Hodgkins lymphoma, a disease entity with far more inflammatory cells than transformed tumor cells. Understanding the biology of individual cells in the complex TME has contributed substantially to shaping models of tumorigenesis. Our studies extend these approaches to evaluate *in vivo* metabolic features of tumor cell types and show that individual cell populations have distinct nutrient uptake programs that may play an important role in therapy response or resistance.

## Methods

### Patient samples

Fresh histology-confirmed clear cell renal cell carcinoma (ccRCC) tumors and matched normal tissue were surgically removed from 14 patients. Tumor and matched normal kidney were processed by mechanical dissociation (human tumor setting two on Miltenyi gentleMACS™) in HBSS with calcium chloride and magnesium chloride. Mechanical dissociation was followed by enzymatic digestion in 435U/mL deoxyribonuclease I (Sigma-Aldrich, D5025) and 218U/mL collagenase (Sigma-Aldrich, C2674) in RPMI supplemented with 10% FBS, 1% glutamine, 1% pen/strep, 1% Hepes, and 0.1% 2-Mercaptoethanol for 30-45min, depending on tissue toughness, at room temperature with 17 rpm agitation. Tissue digests were washed with HBSS without calcium chloride, magnesium chloride, or magnesium sulfate and then incubated in 5mM EDTA for 20min at room temperature with 17rpm agitation. Tumor and matched normal kidney digests were washed with HBSS with calcium chloride and magnesium chloride. Then they were passed through a 70μm filter and ACK-lysed. Patient peripheral blood mononuclear cells (PBMC) were isolated by density gradient centrifugation using Ficoll-Paque (GE Healthcare, 17144002) in SepMate-50 tubes (Stemcell Technologies, 85450) and subsequently ACK-lysed. Single cell suspensions were frozen in 90% FBS 10% DMSO. Batched tumor and matched PBMC were thawed, rested for 10min at 37°C, counted, stained, and analyzed for flow cytometry. All studies were conducted in accordance with the Declaration of Helsinki principles under a protocol approved by the Vanderbilt University Medical Center (VUMC) Institutional Review Board (protocol #151549). Informed consent was received from all patients prior to inclusion in the study.

### Interstitial Fluid Collection & LC/MS metabolite analysis

Tissue interstitial fluid (TIF) was collected from freshly resected ccRCC tumor and matched normal kidney tissue. Specimens were centrifuged against a 0.22μm nylon filter (Corning CLS8169) at 4°C for 5 minutes at 300g. Flow-through TIF was flash-frozen and stored at −80°C prior to batch analysis. Liquid chromatography/mass spectrometry (LC/MS) quantitation of metabolites was performed as described previously^1^.

### Mice

Wild-type C57BL/6J (000664) and BALB/cJ (000651) mice were obtained from the Jackson Laboratory. All mouse procedures were performed under Institutional Animal Care and Use Committee (IACUC)-approved protocols from Vanderbilt University Medical Center and conformed to all relevant regulatory standards. Mice were housed in ventilated cages with at most 5 animals per cage and provided ad libitum food and water. For injectable tumor models, both male and female mice were used. V9302 treatments were administered intraperitoneally twice daily for five days at 25mg/kg for FDG uptake or once at 75mg/kg prior to ^18^F-Gln injection.

### Cell lines

MC38, Renca and CT26 cell lines were grown to 70-90% confluency in DMEM supplemented with 10% FBS or RPMI 1640 supplemented with 10% FBS, 4mM glutamine, 25mM HEPES, essential amino acids, and sodium pyruvate for Renca. 1×10^6^ cells were trypsinized, washed twice in PBS, and injected subcutaneously in 100-200μL of PBS on mouse flanks. Subcutaneous tumors grew for 14 days prior to analysis. cell lines were regularly tested for mycoplasma. The MC38-EL-Thy1.1 cells were generated using a transposon based engineering approach with plasmids that were described previously^2^. MC38 cells were electroporated using the NEON transfection system (ThermoFisher) according to manufacturer’s recommendations for adherent cell lines. 5 million MC38 cells were suspended in electroporation buffer containing 5μg of the plasmid pCMV-M7PB and 15μg of the plasmid pT-EL-thy1.1, which is a bicistronic transposon vector driving expression of an enhanced firefly luciferase as well as Thy1.1 antigen. Cells were magnetically sorted based on expression of Thy1.1 using magnetic beads (Miltenyi 130-121-273).

### Orthotopic renal implantation

For intrarenal Renca injections, survival mouse surgery was performed according to a method previously described^3^. Briefly, mice were anesthetized by isoflurane inhalation at 2-3% and placed on a warming recirculating water pad set at 37°C to maintain body temperature. Using sterile surgical techniques, a 1-cm incision was made in the skin running parallel to the spine, slightly below the ribcage on the right flank. Next, a 1-cm incision was made in the muscle layer in the same location. Using gentle pressure on the mouse abdomen, the right kidney was exteriorized. 5×10^4^ cells resuspended in 100μL of PBS were injected using a 29-gauge needle inserted through the renal capsule into the cortical space. The injection site was swabbed using sterile gauze and the kidney was returned to the body cavity. The abdominal wall was closed using 6-0 monofilament absorbable sutures (AD Surgical; S-D618R13), and the skin was closed using wound clips. Analgesic was provided pre-surgery and 24 hours post-surgery in the form of ketoprofen injections at 5 mg/kg. Wound clips were removed 7 days following the surgery.

### GEMM Tumor Models

PyMT GEMM mice were bred by crossing male transgenic mice expressing the polyoma virus middle T antigen (PyMT) oncoprotein under the MMTV-LTR (Jackson Laboratory 022974) with wildtype females on a similar B6/FVB mixed background. The GEMM mice were from a colony in which all mice expressed two *Vhl* alleles in which exon 1 is flanked by *loxP* sites (Jackson Laboratory 012933), but did not express a Cre transgene and were thus effectively wildtype. Once weaned, female mice were palpated twice a week and tumors were measured in three dimensions with digital calipers. Mice were collected when any tumor had grown to a size of 1 cm in diameter in any dimension, around 5 months of age. Virgin female littermate controls were used in these studies.

The AOM/DSS inflammatory colorectal cancer model was used as previously described^4,5^. In brief, bedding was mixed to normalize microbiome two weeks prior to experimental initiation. Mice were intraperitoneally injected with 12.5 mg/kg AOM and exposed to three 4-day cycles of 3% to 4% DSS (TdB Labs 9011-18-1). Each DSS cycle was followed by a 16-day recovery period. Prior to sacrifice, colonoscopy was performed to confirm tumor development. Mice were weighed every other day throughout the experiment. To allow for larger tumors, mice were euthanized 55 days after completing the last cycle of DSS. Colons were dissected, and tumor tissue was isolated from the mucosa.

### PET-CT Imaging

For individual studies, a group of MC38 tumor-bearing mice were food-restricted overnight. Then the mice received a retro-orbital injection of ~ 37 MBq/0.1 mL of ^18^F-FDG and were returned to their cages. Forty minutes later, the mice were anesthetized under 2% isofluorane and imaged an Inveon microPET (Siemens Preclinical, Knoxville TN) for 20 min. Data from all possible lines of response (LOR) were saved in the list mode raw data format. The raw data was then binned into 3D sinograms with a span of 3 and ring difference of 47. The images were reconstructed into transaxial slices (128 x 128 x 159) with voxel sizes of 0.0815 x 0.0815 x 0.0796 cm^3^, using the MAP algorithm with 16 subsets, 4 iterations, and a beta of 0.0468. For anatomical co-registration, immediately following the PET scans, the mice received a CT scan in a NanoSPECT/CT (Mediso, Washington DC) at an x-ray beam intensity of 90 mAs and x-ray peak voltage of 45 kVp. The CT images were reconstructed into 170 x 170 x 186 voxels at a voxel size of 0.4 x 0.4 x 0.4 mm^3^. The PET-CT images were uploaded into Amide (www.sourceforge.com) and volumetric regions-of-interest were drawn around the tumors. The PET images were normalized to the injected dose and the mean radiotracer concentration within the ROIs were determined.

### ^18^F-FDG and ^18^F-Gln nutrient uptake assay

Tumor-bearing mice were retro-orbitally injected with 1mCi of FDG or ^18^F-Gln synthesized at VUMC^6^. During radiotracer uptake, mice were conscious and had access to food and water. Mice were euthanized humanely, and spleen and tumors were harvested 40min after radiotracer administration. Single cell suspensions of splenocytes were prepared by physical dissociation followed by ACK-lysis. Tumors were chopped, mechanically dissociated on the Miltenyi gentleMACS™ Octo Dissociator with Heaters (setting implant tumor one), and digested in 435U/mL deoxyribonuclease I (Sigma-Aldrich, D5025) and 218U/mL collagenase (Sigma-Alrich, C2674) at 37°C for 30min. After enzyme treatment, tumors were passed through a 70μm filter and ACK-lysed. Cells were resuspended in MACS buffer (PBS +2% FBS +2mM EDTA) and counted using trypan blue with the TC20™ Automated Cell Counter (Bio-Rad). In some cases, tumors from different mice were pooled to achieve higher tumor cell number prior to fractionation to ensure sufficient ^18^F signal. Next, tumor cell suspensions were fractionated using serial magnetic bead positive selection according to the manufacturer’s instructions (all Miltenyi mouse kits: CD45 TIL 130-110-618, EPCAM 130-105-958, Thy1.1 130-121-273, CD4/8 TIL 130-116-480, CD11b 130-049-601, F4/80 130-110-443). Briefly, cells were resuspended at 10 million total cells/90μL MACS buffer and 10μL microbeads for 15min. Then, cell suspensions were applied to LS columns (Miltenyi 130-042-401) in Miltenyi QuadroMACS™ Separators, washed, and eluted according to manufacturer’s instructions. Fractions were resuspended in 1mL of media; 10μL were used for trypan blue staining and TC20 cell count, ~50μL were stained for flow cytometry determination of fraction cellular composition, and 900μL were transferred into 5mL tubes to measure radioactivity. 900μL of 2mL splenocyte suspensions and 5 million total cells from the unfractionated whole tumor were also assayed for radioactivity. The Hidex Automatic Gamma Counter was used with 1min read times to measure time-normalized ^18^F counts per minute (CPM) for each sample. To determine per cell ^18^F-nutrient avidity, time-normalized CPM was divided by the number of viable cells as determined by trypan count. Harvested tissues and cell fractions were kept on ice or at 4°C in RPMI 1640 supplemented with 10% FBS except when noted.

### Flow Cytometry

Single cell suspensions obtained from tumors and spleens were incubated in Fc block (BD 553142) for 10min at room temp, stained for surface markers for 15min at room temp, washed with FACS buffer (PBS +2% FBS) once, and resuspended in FACS buffer for analysis on a Miltenyi MACSQuant Analyzer 10 or 16. For intracellular staining, the eBioscience™ Foxp3/transcription factor staining buffer kit (Fisher 00-5523-00) was used. Surface staining was performed as described above, cells were fix/permed for 20min at 4°C, and then stained for intracellular markers for 30min at 4°C. Ghost Dye Red 780 viability dye (Cell Signaling 18452S) was used identically to surface antibodies. The anti-mouse and cross-reactive antibodies used were: CD45 BV510 (40-F11, Biolegend 103138), B220 e450 (RA3-6B2, ThermoFisher 48-0452-82), CD11b e450 (M1/70, ThermoFisher 48-0112-82), CD11b FITC (M1/70, Biolegend 101206), CD8a AF488 (53-6.7, Biolegend 100723), Ly6C FITC (HK1.4, Biolegend 128006), CD11c PE (N418, BioLegend 117308), FOXP3 PE (FJK-16s, ThermoFisher 12-5773-82), pS6 Ser235/236 PE (D57.2.2E, Cell Signaling 5316S), CD4 PerCP-Cy5.5 (RM4-5, BioLegend 100540), Ly6G PerCP-Cy5.5 (1A8, BioLegend 127616), F4/80 PE-Cy7 (BM8, BioLegend 123114), NKp46 PE-Cy7 (29A1.4, BioLegend 137618), CD3 PE-Cy7 (17A2, BioLegend 100220), CD3 FITC (17A2, BioLegend 100204), CD3 APC (17A2, BioLegend 100236), CD206 APC (C068C2, BioLegend 141708), GLUT1 AF647 (EPR3915, Abcam ab195020), EPCAM PE (G8.8, BioLegend 118206), Thy1.1 PerCP-Cy5.5 (HIS51, ThermoFisher 45-090082), CD45 PE (30-F11, ThermoFisher 12-0451-83), Ly6C BV570 (HK1.4, BioLegend 128030), CD68 BV605 (FA-11, BioLegend 137021). The antihuman antibodies used were: CD45 BV421 (HI30, BioLegend 304032), CD3 APC (UCHT1, BioLegend 300439), CD11b PerCP-Cy5.5 (ICRF44, BioLegend 301328), CD14 BV510 (M5E2, BioLegend 301842), CA9 AF647 (303123, R&D Systems FAB2188R-100UG), Human Fc Block (BD 564220). Flow cytometry data were analyzed using FlowJo v10.7.1.

### Microscopy

MC38 anti-F4/80 microbead-fractionated TAM were mounted onto slides using Wescor Cytopro cytocentrifuge and stained with hematoxylin and eosin following manufacturer’s guidelines (Fisher 23-122952). Images were captured under oil immersion (100x objective) using an Olympus BX53 microscope (Olympus Corporation, Center Valley, PA), an Olympus DP73 camera, and Olympus cellSens Standard imaging software.

### Extracellular flux assay

Tumor cell fractions were obtained as described above. Each fraction was plated at 200,000 live cells/well in at least technical quadruplicate on a Cell-Tak-coated plate (Corning 354240) in Agilent Seahorse RPMI 1640 supplemented with 10mM glucose, 1mM sodium pyruvate, and 2mM glutamine. Cells were analyzed on a Seahorse XFe 96 bioanalyzer using the Mitostress assay (Agilent 103015–100) with 1μM oligomycin, 2μM FCCP, and 0.5μM rotenone/antimycin A. Data were analyzed in Agilent Wave software.

### Single cell sorting and mRNA transcript analysis

CD45^+^ and CD45^-^ tumor cell fractions were obtained as described above. Cell fractions were stained for the indicated surface markers and viability dye and sorted on a BD FACSAria III cell sorter. RNA was isolated from tumor cell populations and unstained whole tumor single cell suspensions using the Quick-RNA™ Microprep Kit (Zymo R1050) according to manufacturer’s instructions. RNA transcripts were quantified using the NanoString nCounter Metabolic Pathways Gene Expression Panel (XT-CSO-MMP1-12) according to manufacturer’s instructions. Transcript levels were normalized to internal control housekeeping genes using the NanoString nSolver software. Differentially expressed genes were defined as having minimally an 8-fold difference between at least two different tumor cell populations in both biological replicates. Gene expression heatmap was generated by normalizing transcript count across each gene. Unsupervised hierarchical clustering of genes and samples were performed using Euclidian distance and linkage was based on unweighted pair group method with arithmetic mean, which is a type of bottom-up agglomerative clustering. Heatmap and clustering was performed using Python3.6 as implemented in the seaborn (v10.1) package using the function *clustermap* with default parameter settings.

### Quantification and Statistical Analysis

Graphs and statistical tests were generated using GraphPad Prism 8 unless otherwise noted.

## Acknowledgements

We thank members of the Rathmell labs for their constructive input. We thank the Justin Balko and Young Kim labs for the use of their tumor dissociators. We thank the Center for Small Animal Imaging at the VUIIS for PET-CT imaging support. We thank the VUMC Radiochemistry core for synthesis and handling of radioactive material. This work was supported by F30 CA239367 (MZM), F30 DK120149 (REB), R01 CA217987 (JCR), NIH T32GM007753 (BTD), R01 DK105550 (JCR), AHA 20PRE35080073 (AA), VA Merit 1I01BX001426 (CSW), Crohn’s and Colitis Foundation 623541 (CSW), American Association for Cancer Research (BIR WKR), and T32 GM007347 (MZM, BIR, REB, AA, CSW). The Vanderbilt VANTAGE Core, including Paxton Baker, provided technical assistance for this work. VANTAGE is supported in part by CTSA Grant (5UL1 RR024975-03), the Vanderbilt Ingram Cancer Center (P30 CA68485), the Vanderbilt Vision Center (P30 EY08126), and NIH/NCRR (G20 RR030956). For the engineered MC38 cells, we thank the cell and genome engineering core of the Vanderbilt O’Brien Kidney Center (P30 DK114809). Flow sorting experiments were performed in the VMC Flow Cytometry Shared Resource by David K. Flaherty and Brittany K. Matlock and was supported by the Vanderbilt Ingram Cancer Center (P30 CA68485) and the Vanderbilt Digestive Disease Research Center (P30 DK058404). This work was supported by grant 1S10OD019963-01A1 for the GE TRACERlab FX2 N and Comecer Hotcell, housed in the Vanderbilt University Institute of Imaging Science - Radiochemistry Core to synthesize ^18^F-Gln.The Inveon microPET was funded by NIH 1S10 OD016245. Figure 1C, Figure 4H, and Extended Data 2C were created using Biorender.

## Extended Data Figure Legend

**Extended Data Fig. 1.**
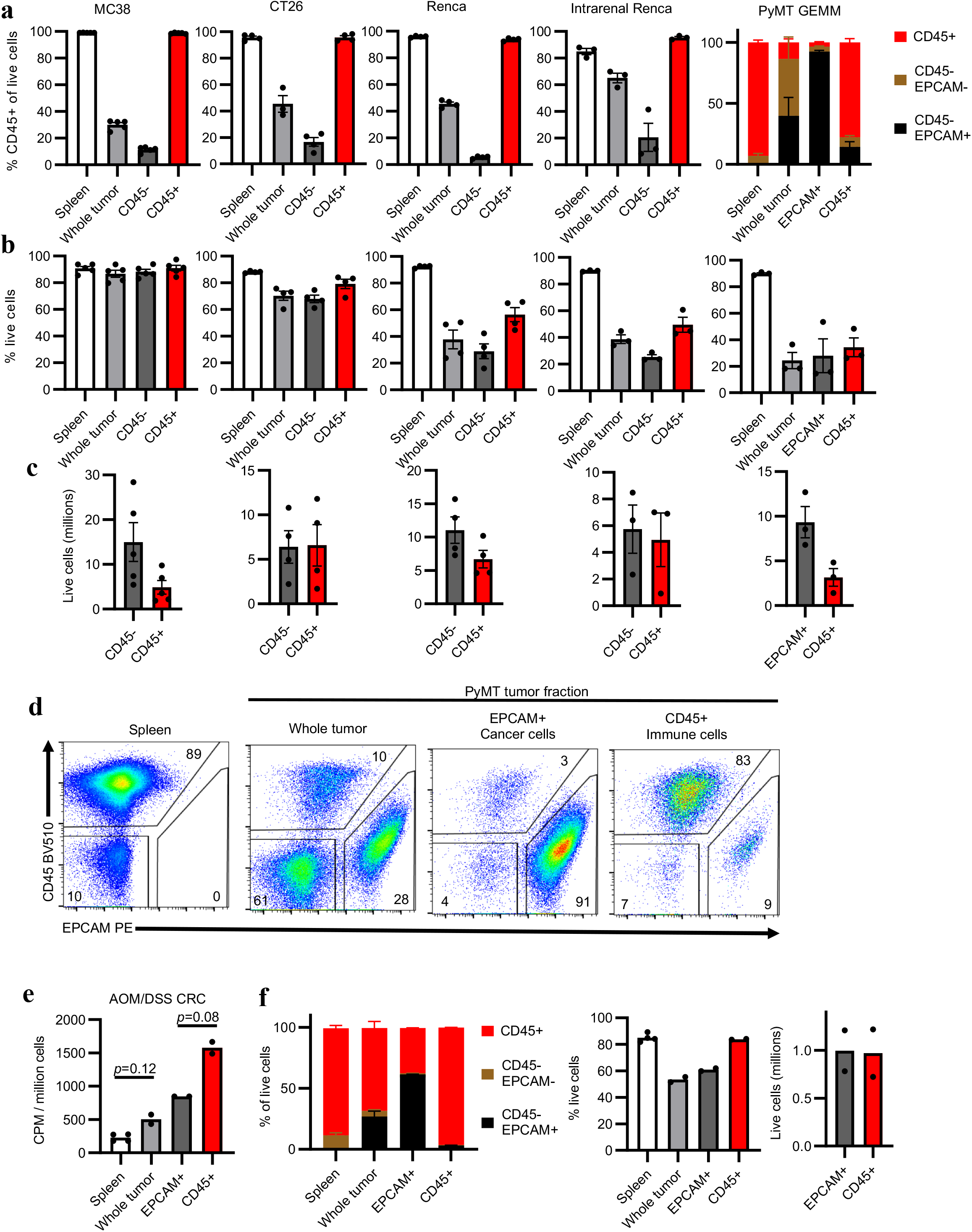
Supporting data for Fig. 1. a-c. Proportion of CD45+ cells (**a**), cell viability (**b**), and live cell yield (**c**) in tumor cell fractions from subcutaneous MC38 (n=5 mice), CT26 (n=4 mice), and Renca (n=4 mice) tumors; intrarenal Renca tumors (n=3 mice); and spontaneous PyMT GEMM (n=3 mice) tumors. **d**, Representative flow cytometry analysis of PyMT whole tumor, CD45+ immune cell, and EPCAM+ cancer cell fractions gated on live cells. **e**, Per cell FDG avidity in designated tumor cell fractions from AOM/DSS CRC tumors (n=2 mice). **f**, Proportion of CD45+ cells, cell viability, and live cell yield from (e). Each data point represents a biological replicate and error bars are SEM. (a-d) are data from representative studies performed independently at least twice. P values were calculated using Welch’s 2-tailed t-test. * *p*<0.05, ** *p*<0.01, *** *p*<0.001. AOM/DSS CRC: azoxymethane/dextran sodium sulfate colorectal cancer; CPM: counts per minute; GEMM: genetically engineered mouse model; PyMT: polyoma virus middle T antigen.

**Extended Data Fig. 2.**
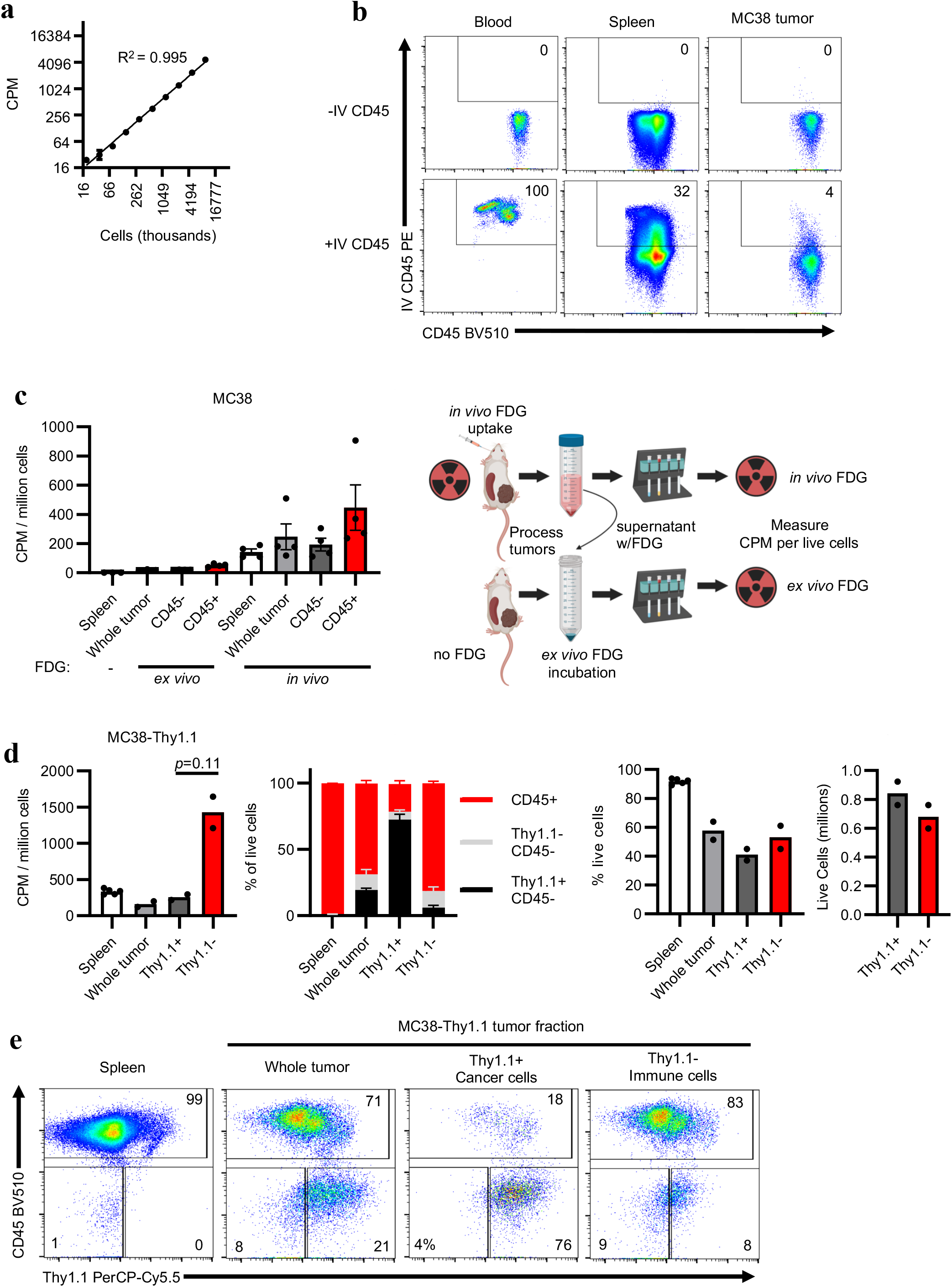
Validation of *in vivo* FDG uptake. **a**, Demonstration of dynamic range of 18F quantification using serially diluted *in vivo* FDG-labelled splenocytes. **b**, Intravenous (IV) anti-CD45 PE staining of leukocytes from designated tissues gated on CD45+ cells. **c**, FDG-hot digest supernatant from *in vivo* labelled MC38 tumors was incubated with FDG-naïve MC38 tumor cells to determine *ex vivo* FDG uptake (n=4 mice). **d**, Per cell FDG avidity in designated tumor cell fractions, proportion of CD45+ and Thy1.1+ cells, cell viability, and live cell yield from MC38-Thy1.1 tumors (n=2 mice). **g**, Representative flow cytometry analysis of MC38-Thy1.1 tumor fractions. Each data point represents a biological replicate and error bars are SEM. (d) are data from a representative study performed independently twice. P values were calculated using Welch’s 2-tailed t-test. * *p*<0.05, ** *p*<0.01, *** *p*<0.001. CPM: counts per minute.

**Extended Data Fig. 3.**
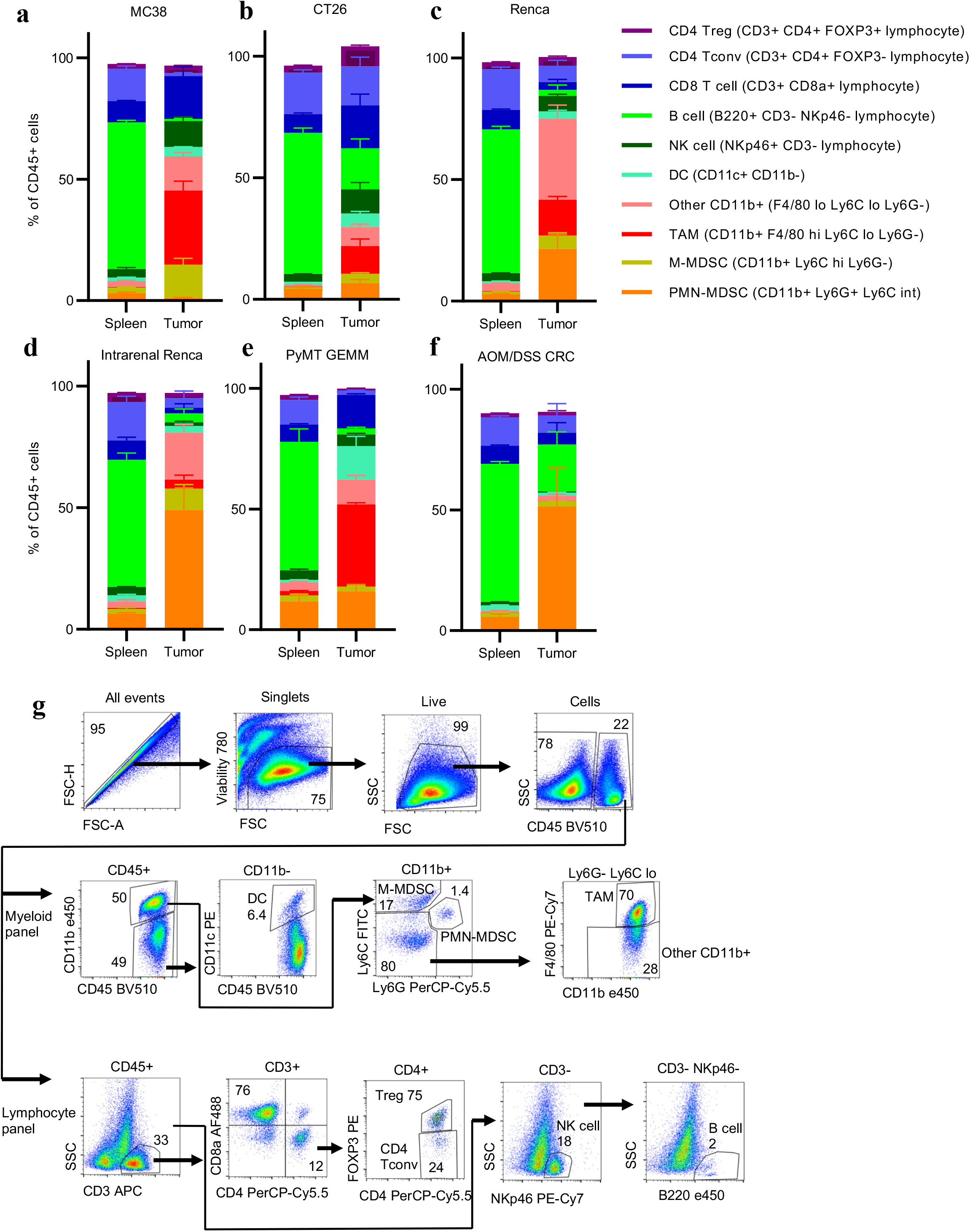
Tumor characterization by flow cytometry. **a-f**, Spleen and tumor immune cell populations from subcutaneous MC38 (n=5 mice), CT26 (n=4 mice), and Renca (n=4 mice) tumors; intrarenal Renca tumors (n=3 mice); spontaneous PyMT GEMM tumors (n=3 mice); and AOM/DSS CRC tumors (n=2 mice). **g**, Gating strategy for immune cell identification using lymphocyte and myeloid-focused antibody panels. Each data point represents a biological replicate and error bars are SEM. Data from a-e are representative of independent experiments performed at least twice. AOM/DSS CRC: azoxymethane/dextran sodium sulfate colorectal cancer; DC: dendritic cell; GEMM: genetically engineered mouse model; M-MDSC: monocytic myeloid-derived suppressor cell; NK cell: natural killer cell; PMN-MDSC: polymorphonuclear myeloid-derived suppressor cell; PyMT: polyoma virus middle T antigen; TAM: tumor-associated macrophage.

**Extended Data Fig. 4.**
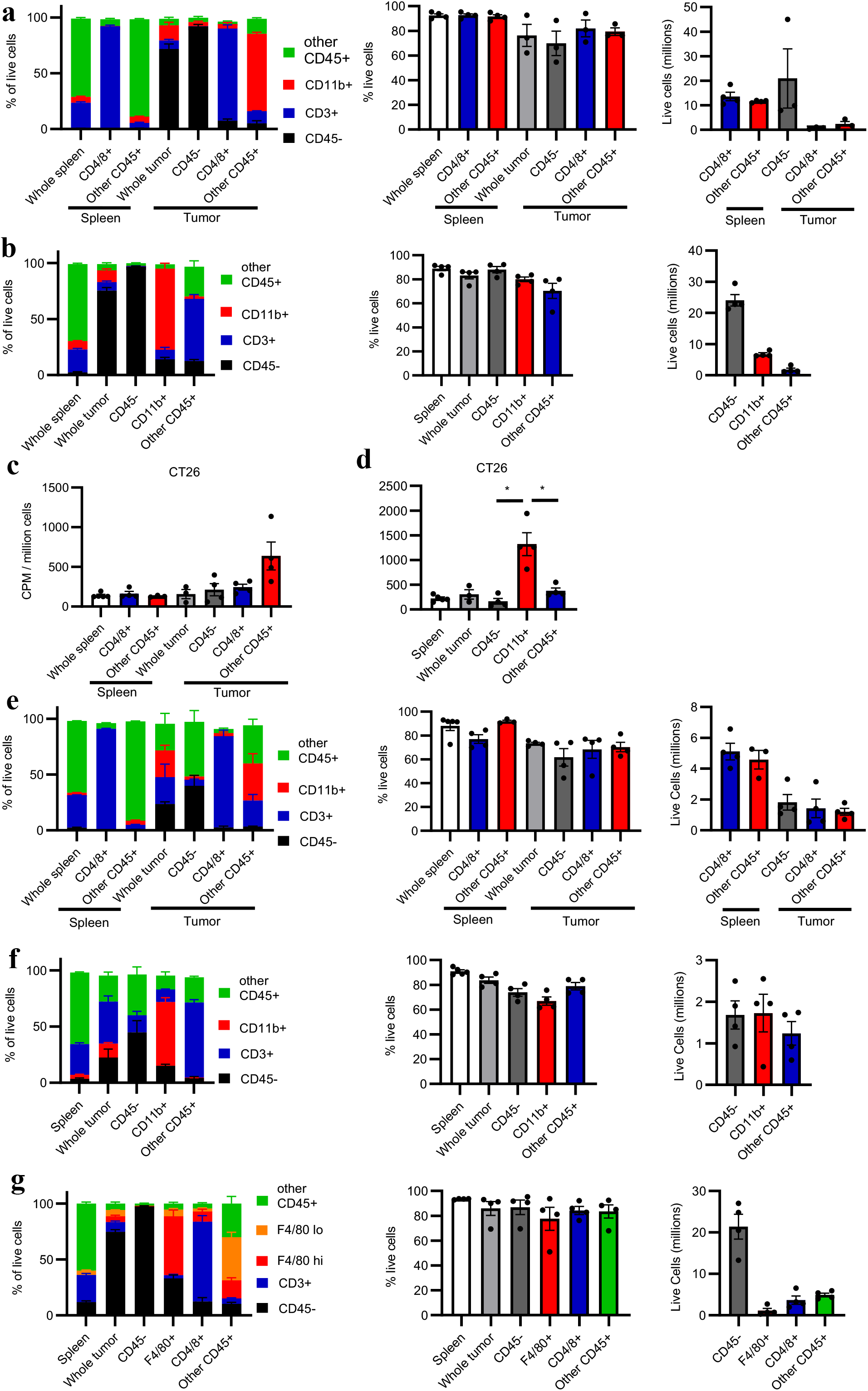
Supporting data for Figure 2. **a-b**, Fraction composition, viability, and live cell yield from MC38 tumor fractions isolated using CD4/8 microbeads (**a**) and CD11b microbeads (**b**) (n=3-4 mice). **c-d**, Per cell FDG avidity in designated CT26 tumor cell fractions using CD4/8 microbeads (**c**) and CD11b microbeads (**d**) (n=4 mice). **e-f**, Fraction composition, viability, and live cell yield from CT26 tumor fractions isolated using CD4/8 microbeads (**e**) and CD11b microbeads (**f**) (n=4 mice). **g**, Fraction composition, viability, and live cell yield from MC38 tumor fractions isolated using F4/80 microbeads. Each data point represents a biological replicate and error bars are SEM. Data are representative of independent experiments performed at least twice. P values were calculated using Welch’s 2-tailed t-test. * *p*<0.05.

**Extended Data Fig. 5.**
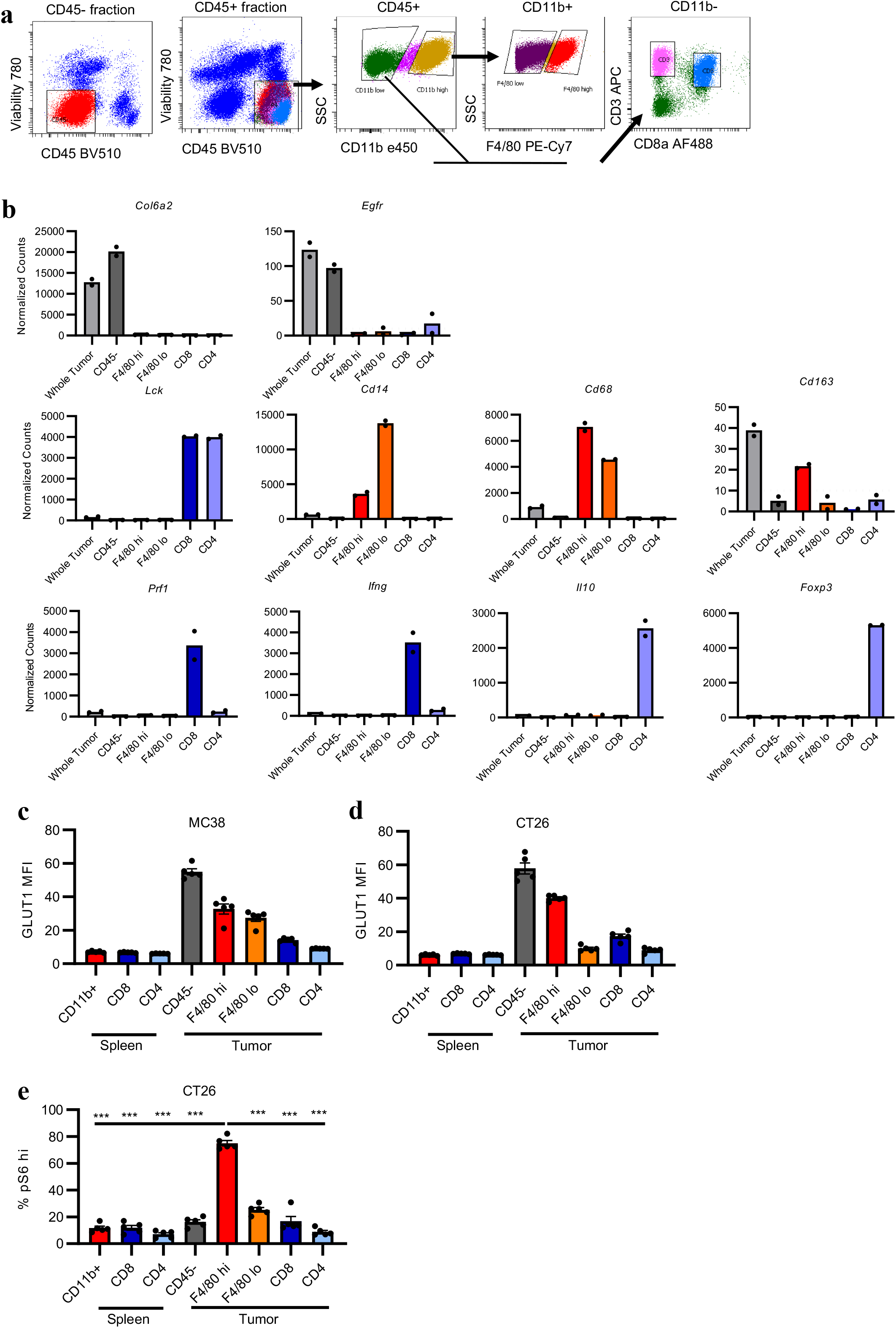
Supporting data for Figure 3. **a**, Sorting gates of MC38 tumor cells used for mRNA transcript analyses. **b**, Expression of selected cell identity markers in flow sorted MC38 tumor cell population. Dotted line approximates limit of detection. **c-d**, GLUT1 levels determined by flow cytometry in MC38 (**c**) and CT26 (**d**) tumor populations. **e,** Quantification of pS6 levels determined by flow cytometry in indicated CT26 tumor and spleen populations. Each data point represents a biological replicate and error bars are SEM. c-e are representative of independent experiments performed at least twice. P values were calculated using the Brown-Forsythe ANOVA test. *** *p*<0.001.

**Extended Data Fig. 6.**
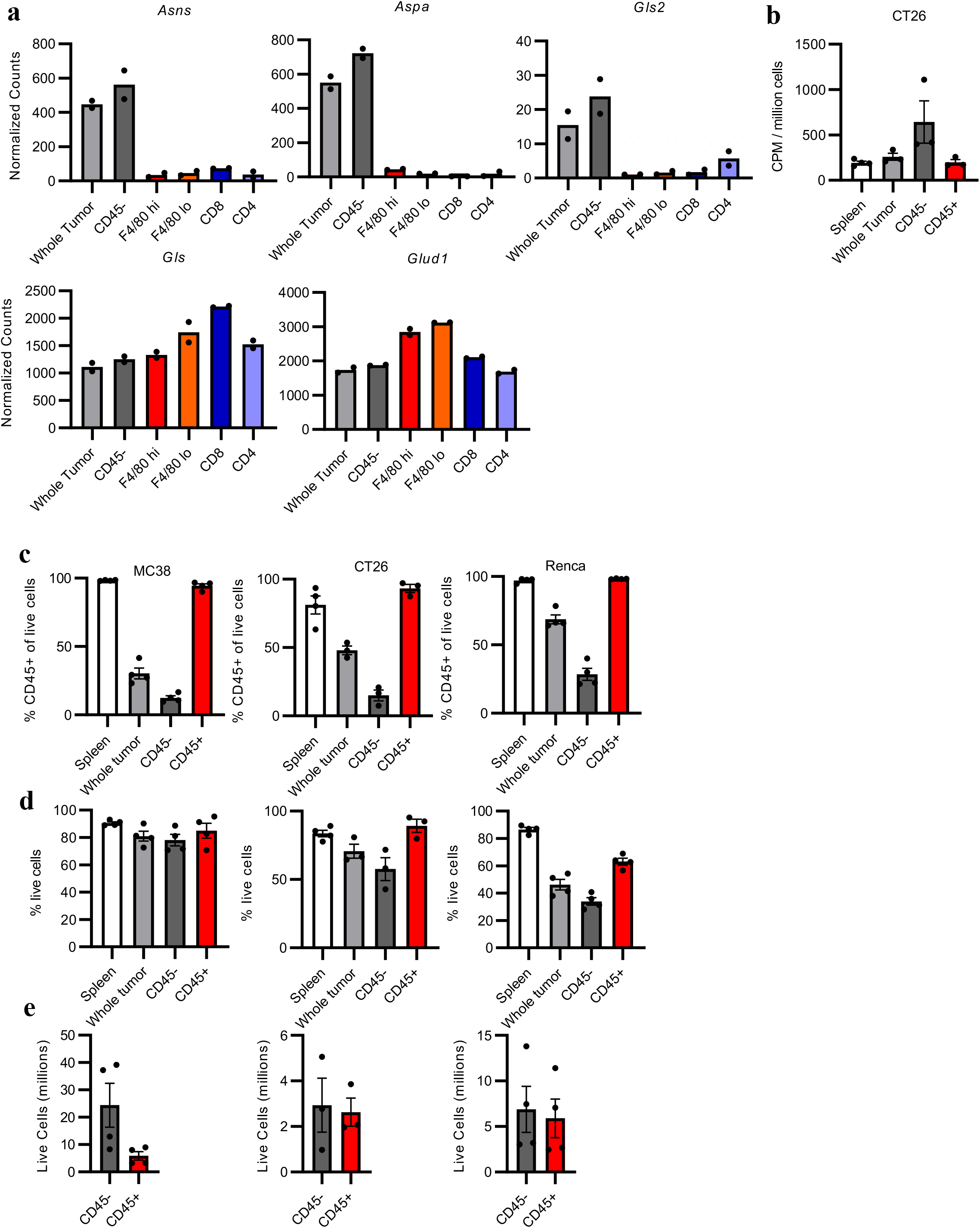
Supporting data for Figure 4a-c. **a**, Amino acid metabolism enzyme mRNA transcript levels of indicated MC38 tumor populations (n=2 mice). Dotted line approximates limit of detection. **b**, Per cell 18F-Gln avidity in designated tumor cell fractions in subcutaneous CT26 tumors (n=3 mice). **c-e**, Proportion of CD45+ cells (**c**), cell viability (**d**), and live cell yield (**e**) in tumor cell fractions from subcutaneous MC38 (n=3-4 mice), CT26 (n=3 mice), and Renca (n=4 mice) tumors. Each data point represents a biological replicate and error bars are SEM.

**Extended Data Fig. 7.**
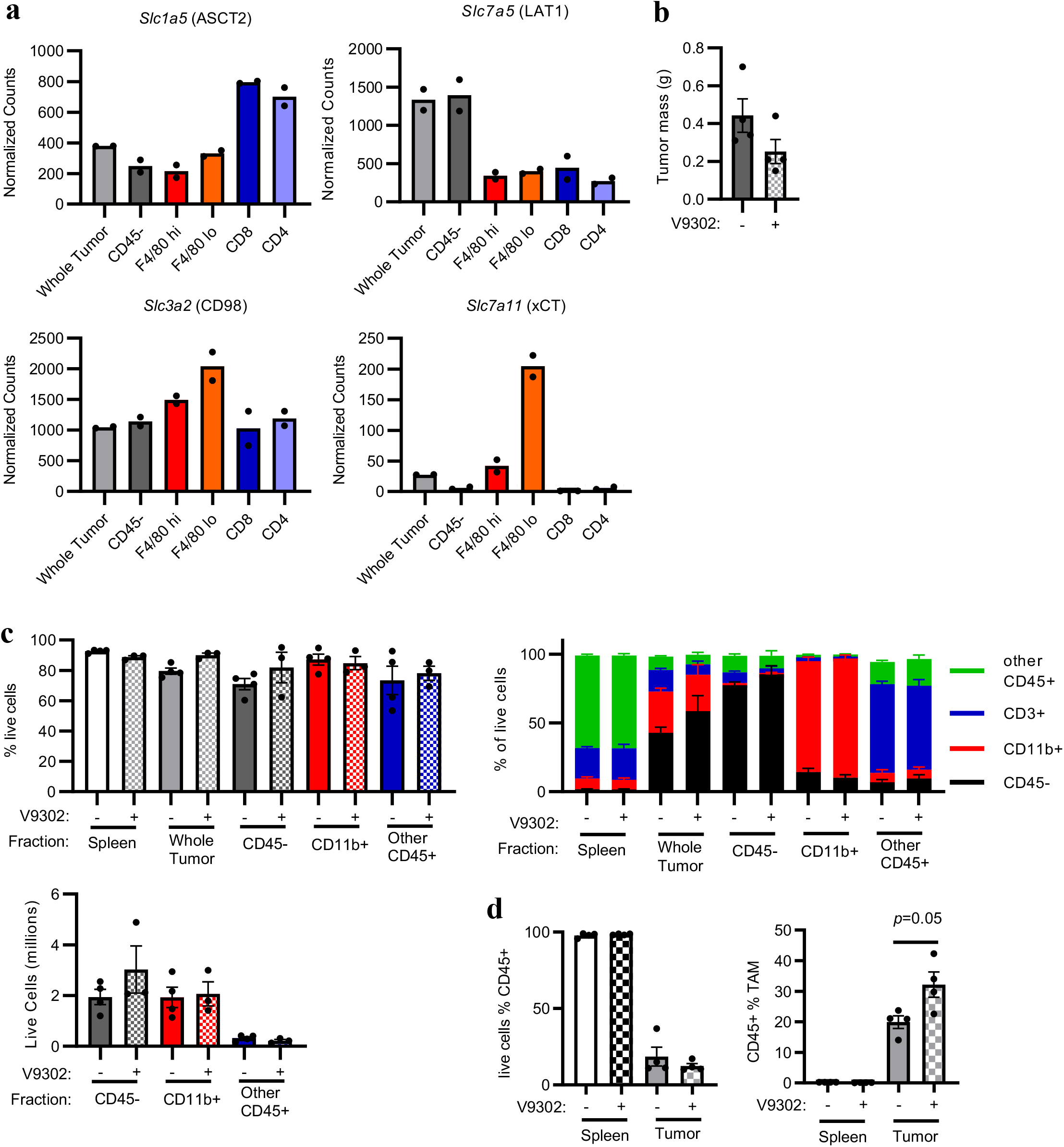
Supporting data for Figure 4d-f. **a**, Glutamine transporter mRNA transcript levels from indicated MC38 tumor populations (n=2 mice). **b,** MC38 tumor mass of DMSO or V9302-treated mice (n=4 mice). **c**, Cell viability, live cell yield, and composition of MC38 tumor fractions from mice treated with DMSO or V9302 (n=3-4). **d,** Flow cytometry analysis of MC38 tumor immune infiltrate from mice treated with DMSO or V9302 (n=4 mice). Each data point represents a biological replicate and error bars are SEM. Data are representative of at least two independent experiments. P values were calculated using Welch’s 2-tailed t-test. * *p*<0.05, ** *p*<0.01, *** *p*<0.001.

## References

1. Chevrier, S. et al. An Immune Atlas of Clear Cell Renal Cell Carcinoma. Cell 169, 736-749 (2017).

2. Slyper, M. et al. A single-cell and single-nucleus RNA-Seq toolbox for fresh and frozen human tumors. Nat. Med. 26, 792–802 (2020).

3. Vander Heiden, M.G., Cantley, L.C. & Thompson, C.B. Understanding the Warburg effect: the metabolic requirements of cell proliferation. Science 324, 1029–1033 (2009).

4. Warburg, O. On the Origin of Cancer Cells. Science 123, 309–314 (1956).

5. Andrejeva, G. & Rathmell, J. C. Similarities and Distinctions of Cancer and Immune Metabolism in Inflammation and Tumors. Cell Metab. 26, 49–70 (2017).

6. Freemerman A.J. et al. Metabolic reprogramming of macrophages: glucose transporter 1 (GLUT1)-mediated glucose metabolism drives a proinflammatory phenotype. J. Biol. Chem. 289, 7884–96 (2014).

7. Jacobs, S. R. et al. Glucose Uptake Is Limiting in T Cell Activation and Requires CD28-Mediated Akt-Dependent and Independent Pathways. J. Immunol. 180, 4476–4486 (2008).

8. Ho, P.C. et al. Phosphoenolpyruvate Is a Metabolic Checkpoint of Anti-tumor T Cell Responses. Cell 162, 1217–1228 (2015).

9. Chang, C.H. et al. Metabolic Competition in the Tumor Microenvironment Is a Driver of Cancer Progression. Cell 162, 1229–1241 (2015).

10. Faubert, B. et al. Lactate Metabolism in Human Lung Tumors. Cell 171, 358–371 (2017).

11. Ma, E. H. et al. Metabolic Profiling Using Stable Isotope Tracing Reveals Distinct Patterns of Glucose Utilization by Physiologically Activated CD8^+^ T Cells. Immunity 51, 856–870 (2019).

12. Oh, M.H. et al. Targeting glutamine metabolism enhances tumor specific immunity by modulating suppressive myeloid cells. J. Clin. Invest. 130, 3865–3884 (2020).

13. Leone, R.D. et al. Glutamine blockade induces divergent metabolic programs to overcome tumor immune evasion. Science 366, 1013–1021 (2019).

14. Liu, P.S. et al. α-ketoglutarate orchestrates macrophage activation through metabolic and epigenetic reprogramming. Nat. Immunol. 18, 985–994 (2017).

15. Johnson, M.O. et al. Distinct Regulation of Th17 and Th1 Cell Differentiation by Glutaminase Dependent Metabolism. Cell 175, 1780–1795 (2018).

16. Sullivan, M.R. et al. Quantification of microenvironmental metabolites in murine cancers reveals determinants of tumor nutrient availability. Elife doi:10.7554/eLife.44235 (2019).

17. Cortese, N. et al. Metabolome of Pancreatic Juice Delineates Distinct Clinical Profiles of Pancreatic Cancer and Reveals a Link between Glucose Metabolism and PD-1+ Cells. Cancer Immunol. Res. 8, 493–505 (2020).

18. Siska, P.J. et al. Mitochondrial dysregulation and glycolytic insufficiency functionally impair CD8 T cells infiltrating human renal cell carcinoma. JCI Insight 2, e93411 (2017).

19. Psychogios, N. et al. The human serum metabolome. PLoS One 6, e16957 (2011).

20. Macintyre, A.N. et al. The glucose transporter Glut1 is selectively essential for CD4 T cell activation and effector function. Cell Metab. 20, 61–72 (2014).

21. Olenchock, B.A., Rathmell, J.C. & Vander Heiden, M.G. Biochemical Underpinnings of Immune Cell Metabolic Phenotypes. Immunity 46, 703–713 (2017).

22. Rathmell, J.C., Elstrom, R.L., Cinalli, R.M. & Thompson, C.B. Activated Akt promotes increased resting T cell size, CD28-independent T cell growth, and development of autoimmunity and lymphoma. European Journal of Immunology 33, 2223–2232 (2003).

23. Pathria, P., Louis, T.L. & Varner, J.A. Targeting Tumor-Associated Macrophages in Cancer. Trends in Immunology 40, 310–327 (2019).

24. Mabuchi, S. et al. Pretreatment tumor-related leukocytosis misleads positron emission tomography-computed tomography during lymph node staging in gynecological malignancies. Nat. Commun. 11, 1364 (2020).

25. Altman, B. J., Stine, Z. E. & Dang, C. V. From Krebs to clinic: Glutamine metabolism to cancer therapy. Nat. Rev. Cancer 14, 749 (2016).

26. Zhang, J., Pavlova, N. N. & Thompson, C. B. Cancer cell metabolism: the essential role of the nonessential amino acid, glutamine. EMBO J. 36, 36:1302–1315 (2017).

27. van Geldermalsen, M. et al. ASCT2/SLC1A5 controls glutamine uptake and tumour growth in triple-negative basal-like breast cancer. Oncogene 35, 3201–3208 (2016).

28. Pathria, G. et al. Targeting the Warburg effect via LDHA inhibition engages ATF 4 signaling for cancer cell survival. EMBO J. 37, e99735 (2018).

29. Qing, G.et al. ATF4 Regulates MYC-Mediated Neuroblastoma Cell Death upon Glutamine Deprivation. Cancer Cell 22, 631–644 (2012).

30. Wang, T. et al. MYCN drives glutaminolysis in neuroblastoma and confers sensitivity to an ROS augmenting agent article. Cell Death Dis. 9, 220 (2018).

31. Zhang, J. et al. Asparagine plays a critical role in regulating cellular adaptation to glutamine depletion. Mol. Cell 56, 205–218 (2014).

32. Gwinn, D.M. et al. Oncogenic KRAS Regulates Amino Acid Homeostasis and Asparagine Biosynthesis via ATF4 and Alters Sensitivity to L-Asparaginase. Cancer Cell 33, 91–107 (2018).

33. Wynn, M.L. et al. RhoC GTPase is a potent regulator of glutamine metabolism and N acetylaspartate production in inflammatory breast cancer cells. J. Biol. Chem. 291, 13715–13729 (2016).

34. Xiao, D. et al. Myc promotes glutaminolysis in human neuroblastoma through directactivation of glutaminase 2. Oncotarget 6, 40655–40666 (2015).

35. Schulte, M.L. et al. Pharmacological blockade of ASCT2-dependent glutamine transportleads to antitumor efficacy in preclinical models. Nat. Med. 24, 194–202 (2018).

36. Bröer, A., Fairweather, S. & Bröer, S. Disruption of amino acid homeostasis by novel ASCT2 inhibitors involves multiple targets. Front. Pharmacol. doi:10.3389/fphar.2018.00785 (2018).

37. Palmieri, E.M. et al. Pharmacologic or Genetic Targeting of Glutamine SynthetaseSkews Macrophages toward an M1-like Phenotype and Inhibits Tumor Metastasis. Cell Rep. 20, 1654–1666 (2017).

38. Lau, A. N. et al. Dissecting cell type-specific metabolism in pancreatic ductal adenocarcinoma. Elife 9, e56782 (2020).

39. Kilgour, M.K. et al. 1-Methylnicotinamide is an immune regulatory metabolite in human ovarian cancer. bioRxiv doi: 10.1101/2020.05.05.077990 (2020).

40. Nair-Gill, E. et al. PET probes for distinct metabolic pathways have different cell specificities during immune responses in mice. J. Clin. Invest. 120, 2005–15 (2010).

41. Halbrook, C. J. et al. Macrophage-Released Pyrimidines Inhibit Gemcitabine Therapy in Pancreatic Cancer. Cell Metab. 29, 1390–1399 (2019).

42. Jeong, H. et al. Tumor-associated macrophages enhance tumor hypoxia and aerobic glycolysis. Cancer Res. 79, 795–806 (2019).

43. Cascone, T. et al. Increased Tumor Glycolysis Characterizes Immune Resistance to Adoptive T Cell Therapy. Cell Metab. 27, 977–987 (2018).

44. Najjar, Y.G. et al. Tumor cell oxidative metabolism as a barrier to PD-1 blockade immunotherapy in melanoma. JCI Insight 4, e124989 (2019).

45. Li, W. et al. Aerobic Glycolysis Controls Myeloid-Derived Suppressor Cells and Tumor Immunity via a Specific CEBPB Isoform in Triple-Negative Breast Cancer. Cell Metab. 28, 87–103 (2018).

46. Chafe, S.C. et al. Carbonic anhydrase IX promotes myeloid-derived suppressor cell mobilization and establishment of a metastatic niche by stimulating G-CSF production. Cancer Res. 75, 996–1008 (2015).

47. Chafe, S.C. et al. Targeting hypoxia-induced carbonic anhydrase IX enhances immune-checkpoint blockade locally and systemically. Cancer Immunol. Res. 7, 1064–1078 (2019).

48. Wenes, M. et al. Macrophage Metabolism Controls Tumor Blood Vessel Morphogenesis and Metastasis. Cell Metab. 24, 701–1714 (2016)

49. Kernstine, K.H. et al. Does Tumor FDG-PET Avidity Represent Enhanced Glycolytic Metabolism in Non-Small Cell Lung Cancer? Ann. Thorac. Surg. 109, 1019–1025 (2020).

50. Kubota, R. et al. Intratumoral distribution of fluorine-18-fluorodeoxyglucose in vivo: high accumulation in macrophages and granulation tissues studied by microautoradiography. J. Nucl. Med. 33, 1972–80 (1992).

## Methods References

1. Sullivan, M. R. et al. Quantification of microenvironmental metabolites in murine cancers reveals determinants of tumor nutrient availability. Elife, doi:10.7554/eLife.44235 (2019).

2. O’Neil, R. T. et al. Transposon-modified antigen-specific T lymphocytes for sustained therapeutic protein delivery in vivo. Nat. Commun. 9, 1325 (2018).

3. Tracz, A., Mastri, M., Lee, C.R., Pili, R. & Ebos, J.M.L. Modeling spontaneous metastatic renal cell carcinoma (mRCC) in mice following nephrectomy. J. Vis. Exp. 86, 51485 (2014).

4. Parang, B., Barrett, C. W. & Williams, C. S. AOM/DSS Model of Colitis-Associated Cancer. Methods Mol. Biol. 1422, 297–307 (2016).

5. Becker, C. et al. In vivo imaging of colitis and colon cancer development in mice using high resolution chromoendoscopy. Gut 54, 950–954 (2005).

6. Hassanein, M. et al. Preclinical Evaluation of 4-[18F] Fluoroglutamine PET to Assess ASCT2 Expression in Lung Cancer. Mol. Imaging Biol. 18, 18–23 (2016).

